# PapRIV, a BV-2 microglial cell activating quorum sensing peptide

**DOI:** 10.1101/2020.10.05.327148

**Authors:** Yorick Janssens, Nathan Debunne, Anton De Spiegeleer, Evelien Wynendaele, Marta Planas, Lidia Feliu, Alessandra Quarta, Christel Claes, Debby Van Dam, Peter Paul De Deyn, Peter Ponsaerts, Matthew Blurton-Jones, Bart De Spiegeleer

## Abstract

**Background:** Quorum sensing peptides (QSPs) are bacterial peptides produced by Gram-positive bacteria to communicate with their peers in a cell-density dependent manner. These peptides do not only act as interbacterial communication signals, but can also have effects on the host. Compelling evidence demonstrates the presence of a gut-brain axis and more specifically, the role of the gut microbiota in microglial functioning. The aim of this study is to investigate microglial activating properties of a selected QSP (PapRIV) which is produced by *Bacillus cereus* species.

**Methods:** Gastro-intestinal transport of the peptide is investigated using the *in vitro* Caco-2 model while transport over the blood-brain barrier is investigated in mice using multiple time regression experiments. Microglial activation is assessed using ELISA, fluorometry, immunoblotting, qPCR and phase-contrast microscopy. *In vivo* plasma detection and *ex vivo* metabolization experiments are performed using UHPLC-MS^2^ and UHPLC-UV/MS, respectively.

**Results:** PapRIV showed *in vitro* activating properties of BV-2 microglia cells and was able to cross the *in vitro* Caco-2 cell model and pass the blood-brain barrier *in vivo*. *In vivo* peptide presence was also demonstrated in mouse plasma. The peptide caused induction of IL-6, TNFα and ROS expression and increased the fraction of ameboid BV-2 microglia cells in an NF-κB dependent manner. Different metabolites were identified in serum, of which the main metabolite (DLPFEH) still remained active.

**Conclusions:** PapRIV is thus able to cross the gastro-intestinal tract and the blood-brain barrier and shows *in vitro* activating properties in BV-2 microglia cells, hereby indicating a potential role of this quorum sensing peptide in gut-brain interaction.

## Background

Quorum sensing is a cell-cell communication system used by micro-organisms to sense the density of their peers by secreting quorum sensing molecules. By using this system, gene expression is regulated in response to the microbial cell density. A variety of cell functions such as expression of virulence factors, biofilm formation, competence and sporulation are controlled by this communication system [1]. The process of quorum sensing is not only limited to bacteria, but it is also observed in other microbial cell types such as yeasts and fungi [2, 3]. Different types of quorum sensing molecules in bacteria exist: Gram-negative bacteria mostly use N-acyl homoserine lactones, while Gram-positive bacteria produce oligopeptides for their communication, which are called quorum sensing peptides (QSPs) [4–6]. Other quorum sensing molecules, such as furan borate derivatives and other miscellaneous molecules, exist as well [7, 8]. The QSPs are produced as large pro-peptides, secreted outside the microbial cell by ATP-binding cassette transporters, whilst hydrolyzed to the active QSP. They then interact with neighboring microbial cells via two possible mechanisms: either with membrane-located receptors (mainly histidine kinases) for signal transduction, or directly with cytoplasmic sensors (*e.g*. the RNPP family) after being transported over the microbial cell membrane by oligopeptide permeases [4]. The QSPs are chemically and microbiologically described in the Quorumpeps^®^ database which currently contains over 350 different peptides [9]. Recently, it has been found that these peptides not only influence micro-organisms but can also affect cells of the host. For example, QSPs promote tumor cell invasion and angiogenesis *in vitro*, thereby promoting epithelial-mesenchymal transition and metastasis [10, 11]. The human immune system also makes use of these molecules for controlling infection. Mast cells are able to sense CSP-1, a QSP produced by *Streptococcus pneumoniae*, hereby triggering degranulation and the release of antibacterial mediators [12]. *In vitro* research also suggests that QSPs influence host muscle wasting diseases [13]. QSPs can cross the blood-brain barrier and thus potentially influence brain cells [14]. Indeed, an *in vitro* screening of 85 different QSPs indicated that some peptides have the ability to exert biological effects on different types of brain cells [15].

Microbial dysbiosis is observed in a variety of neurodevelopmental-, neurodegenerative- and psychiatric disorders such as autism spectrum disorders (ASD), schizophrenia, Alzheimer’s disease (AD), major depressive disorder, and Parkinson’s disease (PD) [16–24]. A causal relationship between these gut microbiota and brain disorders is becoming increasingly evident: fecal transfer of human ADHD, PD and AD patients aggravates symptoms in *in vivo* mice models of these disorders [25–27]. Alternatively, transfer of a ‘healthy’ microbiota reduced amyloid and tau pathology in an AD mouse model [28]. This microbiota-brain association is considered part of a bi-directional communication pathway between the gut and brain, called the gut-brain axis [29]. Various communication routes including the immune system, the vagus nerve, the enteric nervous system and microbial metabolites, such as short chain fatty acids, amino acids and peptidoglycans are proposed as mediators of this axis, but many factors remain largely unknown [30]. Peptides also contribute in this microbiome-to-brain signaling as interactions of the microbiota with gut hormones and entero-endocrine peptides are observed [31]. Microglial dysfunction and activation is present in a variety of neuronal conditions such as AD, ASD, multiple sclerosis, PD and amyotrophic lateral sclerosis [25, 32–37]. Germ-free mice display global defects in microglia with altered cell proportion and an immature phenotype, indicating that the gut microbiome plays a role in microglial maturation and functioning during development and aging [32, 38]. When transplanting faeces from human AD or PD patients to *in vivo* mouse models, cognitive and physical impairments are aggravated, which is partly mediated by enhanced microglial activation in the brain [25, 26]. Prebiotics are already being developed that target this gut microbiota-microglial axis for the treatment of AD. Sodium oligomannate, which will shortly be studied as an investigational drug in a large AD phase III global clinical trial, suppresses gut dysbiosis together with the associated phenylalanine/isoleucine accumulation in an AD mouse model, resulting in reduced microglial activation, amyloid-β deposition, tau phosphorylation and amelioration of cognitive impairment [39]. All these studies indicate that the gut microbiota can influence neurodevelopmental-, neurodegenerative- and psychiatric disorders by regulation of microglia cells. However, the exact mechanisms on how the gut microbiota influence these microglial cells remains largely unknown.

PapRIV is a heptapeptide (SDLPFEH) originating from the *Bacillus cereus* group, which comprises a number of highly related species. The *B. cereus* group (sensu lato), which is widespread in soil and food, is generally considered as an opportunistic human pathogen because it triggers food-borne gastroenteritis and some non-gastro-intestinal infections like pneumonia and endophthalmitis, as well as infections resembling to anthrax. However, presence of *B. cereus* in the human gastro-intestinal tract is already been demonstrated without provoking illness, indicating the symbiotic life cycle of this species [40]. The diseases associated with *B. cereus* are caused by several cytotoxic products, which are produced by activation of the PlcR quorum sensing system. The PapRIV peptide is translated as a 48-amino acid polypeptide which is secreted out of the cell under influence of the N-terminal signaling sequence and extracellularly processed by NprB proteases to form the active PapRIV heptapeptide [41, 42]. Instead of binding to a membrane-coupled sensor receptor and activating a two-component signaling system, PapRIV binds directly to its regulatory cytoplasmatic PlcR protein after being imported in the cell by the ABC transporter family member Opp (oligopeptide permease). This binding activates the regulatory PlcR protein resulting in a conformational change, binding to the promotor region and transcriptional activation of PlcR target genes [43, 44]. Activation of this quorum sensing system induces production of extracellular virulence factors, such as enterotoxins, haemolysins, cytotoxins and various degradative enzymes (*e.g*. proteases). However, the effects of the peptide itself towards the host remains unexplored. Here, we demonstrate for the first time microglial activating properties of a QSP, indicating the potential of these peptides as mediators of the gut-brain-microglia axis.

## Methods

### Peptides and Reagents

Synthetic PapRIV was purchased from GL Biochem (Shanghai, China). The alanine scan, metabolites and scrambled control were synthesized using solid-phase peptide synthesis (Supplementary method S1). The quality of all peptides was determined using an in-house developed QC method and a purity of 95% or more was found for each sequence. Calcium dichloride dihydrate, magnesium sulphate, potassium chloride, sodium dihydrogen phosphate hydrate, HEPES, sodium lactate and urethane were purchased from Sigma-Aldrich (Diegem, Belgium), while Bovine Serum Albumin (BSA), sodium iodide, sodium dihydrogen phosphate monohydrate were obtained from Merck KGaA (Darmstadt, Germany). Sodium chloride and disodium hydrogen phosphate dihydrate were obtained from VWR (Leuven, Belgium). Calcium dichloride and D-glucose were purchased from Fluka (Diegem, Belgium) and dextran from AppliChem GmbH (Darmstadt, Germany). Water was purified using an Arium 611 Pro VF purification system (Sartorius, Göttingen, Germany) to laboratory-graded water (18.2 MΩ × cm).

### Animals

Female, Institute for Cancer Research, Caesarean Derived-1 (ICR-CD-1) mice (Envigo, Venray, The Netherlands) of age 7-10 weeks and weighing 26-30 g, were used during the blood-brain barrier (BBB) transport experiments and C57Bl6/J WT mice, aged 3-24 months, were used for detection of PapRIV in plasma. Feed and water were provided ad libitum. All animal experiments were performed in strict accordance with the Belgian legislation RD 31/12/2012 and the Ethical Committee principles of laboratory animal welfare; the protocols were approved by the Ethical Committee of Ghent University, Faculties of Veterinary Medicine and Medicine and Health Sciences (approval numbers 2014-128 and ECD 19-17).

### Cells

BV-2 cells were a kind gift from Prof. Alba Minelli and were grown in RPMI medium supplemented with 10% fetal bovine serum (FBS) and 1% penicillin-streptromycin solution (Life Technologies, Belgium). The cells were cultured in culture flasks (Greiner, Belgium) in an incubator set at 37°C and 5% CO_2_. When confluent, cells were detached using a cell scraper and diluted 1:20 approximately every 4 days. Experiments were performed in serum-free DMEM without phenol red.

SH-SY5Y neuroblast cells (Sigma-Aldrich, Diegem, Belgium) were grown in F12:MEM (50/50 v/v) medium supplemented with 15% FBS, 1% non-essential amino acids (NEAA), 2mM L-Glutamine and 1% penicillin/streptomycin solution (Life Technologies, Belgium) in an incubator set at 37°C and 5% CO_2_. When confluent, cells were detached using 0.25% Trypsin-EDTA.

### Caco-2 assay

Caco-2 cells were grown in DMEM supplemented with 10% FBS, 1% NEAA and 1% penicillin/streptomycin solution. A total of 3 x 10^5^ cells per filter were seeded in a 12-well plate and left for differentiation during 21-29 days (medium change every other day). Monolayer formation was checked by measuring the TEER values which should be above 0.30 kΩ x cm^2^ [45]. After washing with Hank’s balanced salt solution, 400 μL of a 1 μM peptide solution was added to the apical side of the filter. After 30, 60, 90 and 120 minutes, 100 μL of the acceptor compartment was taken for UPLC-MS/MS analysis (Supplementary method S2). Linear curve fitting was used to calculate the apparent permeability coefficient (P_app_), as sink conditions were achieved. The P_app_ was determined from the amount of peptide transported per time unit and calculated using the following equation:

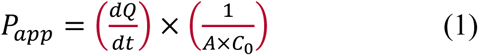

with

dQ/dt = the steady-state flux (pM/s), experimentally obtained
A= the surface area of the filter (= 1.12 cm^2^)
C_0_= the initial concentration in the donor compartment (= 1 x 10^6^ pM)

The reduction in acceptor concentration was also taken into account after every sampling. The cumulative amount in the acceptor compartment is defined as:

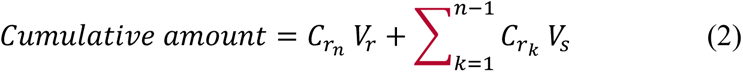

with

*C_r_k__* = concentration at sample point k in acceptor compartment
*V_r_* = volume in the receiver chamber
*V_s_* = volume sampled
n = times sampled

### Detection of PapRIV in mouse plasma

Sixty μL of mouse plasma was used for the detection of PapRIV and prepared using solid-phase extraction (SPE). Mouse plasma was mixed with an ACN/DMSO mixture acidified with FA (94/3/3 V/V). After 30s vortexing and sonification (30s), the sample was boiled for 1min at 100° followed by centrifugation (30s, 10,000g). One hundred μL of the supernatant was then mixed with 800 μL ACN/DMSO mixture (97/3 V/V) acidified with 0.1% FA. This sample was then loaded on a MonoSpin Amide SPE column (GL Sciences Inc., Japan) conditioned with a water/ACN mixture (90/10 V/V) basified with 0.1% NH_4_OH and equilibrated with the same ACN/DMSO mixture. The sample was eluted using a water/ACN/DMSO mixture (75/20/5 V/V) acidified with 0.1% FA. The eluent was brought into vials which were coated with an albumin based anti-adsorption solution, which considerably decreases the adsorption to glass for certain peptides and improves the overall sensitivity of the method [46]. Ten μL of the eluent was injected on an Acquity UPLC^®^ BEH C18 (2.1 x 100 mm; 1.7 μm) column equipped with a guard column. Column temperature was maintained at 60°C. Mobile phase A consisted of water/ACN/DMSO (93/2/5 V/V) + 0.1% FA, Mobile phase B consisted of water/ACN/DMSO (2/93/5 V/V) + 0.1% FA. A gradient ranging from 100% mobile phase A to 60% mobile phase B over 12 minutes was used. MS analysis was performed using a Quadrupole-Time-of-flight system (SYNAPT G2-Si) (Waters, Milford, USA). The SYNPAT G2-Si with electrospray ionisation was set in positive mode. Ionization was conducted using a needle voltage of 3.0 kV, a cone voltage of 20 V and a source temperature of 120°C. Nitrogen was used as sheath and auxiliary gas at a temperature of 500°C. Argon was used as collision gas at an energy of 32 V. Detection was performed using a fixed mass and collision energy on the first quadrupole set on the mother ion (844.35 ± 0.5 *m/z*) and MS/MS acquisition over the 100-1450 *m/z* range using the second TOF analyser. Samples were considered positive when a signal appeared at the expected retention time (Δ < 1%) and when at least three identification ions (one parent ion and two daughter ions) of the peptide were found [47].

### Peptide ^125^I radiolabelling

PapRIV and the controls BSA and dermorphin (Hanhong, Shanghai, China) were iodinated using the Iodo-Gen^®^ method as previously described [48]. Briefly, 0.1 μmol of the lyophilized peptide was dissolved in 100 μL of phosphate buffer (pH 7.4, 25 mM). A Iodo-Gen^®^ coated tube (Thermo Scientific, Merelbeke, Belgium) was first of all rinsed with 1 mL of phosphate buffer. Subsequently, 50 μL of sodium iodide solution (1.1 μmol/mL) and 1 mCi of Na^125^I solution (Perking Elmer, Zaventem, Belgium) were transferred into this Iodo-Gen^®^ coated tube. The oxidation reaction was allowed to proceed for six minutes at room temperature, after which the iodonium solution was transferred to 50 μL of peptide solution (1 mM). The iodination reaction of the peptide was allowed to proceed for another six minutes at room temperature. Finally, the iodinated peptide was purified using a silver filter (Thermo Scientific, Merelbeke, Belgium).

### Multiple time regression analysis

In order to determine whether the peptide could enter the brain, *in vivo* multiple time regression (MTR) analysis was performed. ICR-CD-1 mice were anesthetized intraperitoneally using a 40% urethane solution (3 g/kg). Then, the jugular vein and carotid artery were isolated and 200 μL of the radiolabeled peptide solution, diluted to 30,000 cpm/μL using Lactated Ringer’s solution containing 1% of BSA (LR/BSA), was injected into the jugular vein. At specified time points after injection (i.e. 1, 3, 5, 10, 12.5 and 15 min, with start and end in duplicate), blood was obtained from the carotid artery followed by decapitation of the mouse. The isolated brain was weighed and radioactivity measured in a gamma counter for 5 minutes (Wallac Wizard automatic gamma counter, Perkin Elmer, Shelton, CT, USA), as well as from 50 μL serum, which was obtained by centrifuging the collected blood at 10,000 g for 15 min at 21°C. The linear and biphasic modeling of the multiple time regression analysis was performed as previously described [14].

### Capillary depletion

We performed capillary depletion to determine whether the peptides, taken up by the brain, completely crossed the capillary wall into the tissue rather than just being trapped by and in the endothelium. The method of Triguero *et al*., as modified by Gutierrez *et al*., was used [49, 50]. ICR-CD-1 mice were first anesthetized intraperitoneally using a 40% urethane solution (3 g/kg). After isolation of the jugular vein, 200 μL of the iodinated peptide solution, diluted to 10 000 cpm/μL using LR/BSA, was injected in the jugular vein. Ten minutes after injection, blood was collected from the abdominal aorta and the brain was perfused manually with 20 mL of Lactated Ringer’s buffer after clamping the aorta and severing the jugular veins. Subsequently, the brain was collected, weighed and the radioactivity measured in the gamma counter for 5 minutes. Then, the brain was homogenized with 0.7 mL of ice-cold capillary buffer (10 mM HEPES, 141 mM NaCl, 4 mM KCl, 2.8 mM CaCl_2_, 1 mM MgSO_4_, 1 mM NaH_2_PO_4_ and 10 mM D-glucose adjusted to pH 7.4) in a pyrex glass tube and mixed with 1.7 mL of 26% ice-cold dextran solution in capillary buffer. The resulting solution was weighed and centrifuged in a swinging bucket rotor at 5,400 g for 30 min at 4°C, after measuring the radioactivity in the gamma counter. Pellet (capillaries) and supernatant (parenchyma and fat tissues) were also collected, weighed and measured in a gamma counter. After centrifuging the obtained blood (10,000 g, 21°C, 15 min), the radioactivity of 50 μL serum was measured in a gamma counter as well.

### Brain-to-blood transport

We quantified the amount of peptide exported out of the brain as previously described [51]. ICR-CD-1 mice were anesthetized intraperitoneally using a 40% urethane solution (3 g/kg). Then, the skin of the skull was removed and using a 22 G needle marked with tape at 2 mm, a hole was made into the lateral ventricle at the following coordinates: 1 mm lateral and 0.34 mm posterior to the bregma. The anesthetized mice received an intracerebroventricular (ICV) injection of 1 μL of the diluted iodinated peptide solution using LR/BSA (25,000 cpm/μL) by pumping the peptide solution at a speed of 360 μl/h for 10 s using a syringe pump (KDS100, KR analytical, Cheshire, UK). At specified time points after ICV-injection (i.e. 1, 3, 5, 10, 12.5 and 15 min), blood was collected from the abdominal aorta and subsequently the mouse was decapitated. Then, the whole brain was collected, weighed and measured in a gamma counter for 5 minutes. Fifty μL of serum, which was obtained by centrifuging the collected blood at 10 000 g during 15 min at 21°C, and the background was also measured in a gamma counter. The efflux half-life was calculated from the linear regression of the natural logarithm of the residual radioactivity in brain versus time as follows:

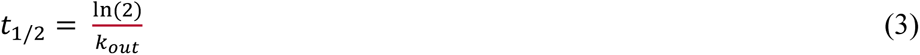

where k_out_ is defined as the efflux rate constant calculated as the negative value of the slope of the linear regression, applying first order kinetics.

### IL-6 and TNFα determination

To investigate the microglia activating properties of the peptide, IL-6 and TNFα levels were determined in cell-free supernatants of BV-2 microglia cells after treatment. BV-2 cells (2 x 10^5^ cells/well) were seeded in 24-well plates and treated with peptide for 20 hours. ELISA was performed according to the supplier’s protocol (eBioscience, Vienna, Austria). Briefly, after incubation with biotinylated detection antibody, avidin-HRP conjugate and subsequently chromogenic tetramethylbenzidine (TMB) substrate were added. Absorbance was measured at 450 nm and 570 nm using the Multiskan Ascent 354 microplate reader (Thermofisher, Waltham, USA). Concentrations were determined using the standard curve generated using known concentrations of TNFα and IL-6.

### qPCR

To confirm the observed ELISA results at mRNA level, a qPCR experiment was performed. After incubation of the BV-2 cells with PapRIV or controls, cells were lysed in RLT buffer (Qiagen, Hilden, Germany) supplemented with 1% β-mercaptoethanol. The lysate was stored at −80°C until RNA extraction. RNA was extracted using the RNeasy Mini Kit (Qiagen, Hilden, Germany). DNAse steps were included in the protocol to remove possible DNA contamination. RNA purity and concentration were assessed using spectrophotometry (NanoDrop), while RNA quality was evaluated using capillary electrophoresis (Fragment Analyzer). After extraction, the RNA was immediately converted to cDNA which was stored at −20°C until qPCR analysis. Ppia and Rer were chosen as suitable control genes based on the GeNorm and Normfinder algorithms [52, 53]. Used primers are given in Table 1. qPCR cycling was performed using a LightCycler 480 (Roche), with Cq values being calculated using the second derivative threshold method.

**Table 1:**
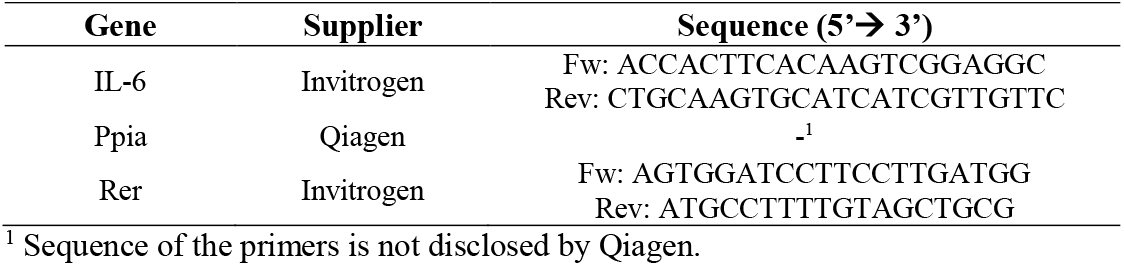
Primers for BV-2 qPCR

### ROS assay

Microglia activation is accompanied by an increased reactive oxygen species (ROS) production. The ROS assay was performed using a fluorometric intracellular ROS kit according to the supplier’s protocol (Sigma-Aldrich, Diegem, Belgium). In brief, 4 x 10^4^ cells/well were seeded in a black 96-well plate. Four hours post seeding, 100 μL of master reaction mix was added to the wells and incubated for 1 h after which cells were treated with PapRIV dissolved in PBS for 24h. Fluorescence intensity was measured using an EnVision fluorometric plate reader at λ_ex_ = 492/λ_em_ = 535 nm (Perkin Elmer, Zaventem, Belgium).

### Morphological analysis

We investigated the morphology since microglia change their morphology from a branched structure to a more round, ameboid structure after activation. BV-2 cells were seeded in 1 x 10^4^ cells/well in a 24-well plate (360 μL); this way direct cell-cell contacts, which have an influence on morphology, are avoided. Four hours post-seeding, cells were treated with 40 μL peptide, placebo (H_2_O), medium or positive control (LPS 1 μg/mL). For the assessment of the morphology, one picture in the center of each well (100x magnification) was taken after twenty hours with an Olympus CKX53 phase-contrast microscope equipped with an XC30 CCD camera (Olympus NV, Antwerp, Belgium). The number of branched and ameboid cells was counted with the cell counter plugin of ImageJ.

### Immunoblotting

To assess nuclear translocation of NF-kB, cells were lysed after 20h using the NE-PER™ protocol (Thermo Scientific). Using this kit, the cytoplasmic and nuclear protein fractions are separated from each other. To asses IκBα expression, cells were lysed with RIPA buffer. Protein content of the lysates was determined using the modified Lowry assay (Thermo Scientific). Proteins (20 μg) were separated using an Any kD gel (SDS-PAGE) and transferred to a nitrocellulose membrane (BioRad, Temse, Belgium). Aspecific binding sites were blocked for 30 min using TBS + 1% casein (BioRad, Temse, Belgium). Western blot was performed using the following primary antibodies overnight (4°C): anti-p65 NFkB (1/500), anti-IκBα (1/1000) and anti-β-actin (1/1000), β-tubulin (1/4000) and anti-Histon H3.3 (1/5000) were used as loading controls. Goat anti-rabbit-AP antibody was used for detection (1/10000) (60 min). All antibodies were purchased at Thermo Scientific (Merelbeke, Belgium). Finally, the BCIP/NBT substrate was added and the results were analyzed using the GelDoc EZ imager and Image Lab software (BioRad, Temse, Belgium). TBS buffer was used for washing between the steps.

### MTT assay

SH-SY5Y cells were seeded at a density of 5 x 10^4^ cells/well in a 96-well plate and incubated for 24 h. Next, medium was removed and replaced by 200 μL conditioned medium of BV-2 cells which were treated for 20 h with PapRIV. After 24 h incubation, 20 μL MTT reagent (12mM) was added and incubated for 3 h. Finally, the medium was removed and replaced by 150 μL DMSO and measured at 570 nm with a microplate reader (Thermofisher, Waltham, USA).

### *Ex vivo* metabolization

Peptide solution (0.1 mg/mL) was incubated with serum, brain, liver, kidney or faeces homogenate at 37°C. After 0, 5, 10, 30 and 60 minutes, aliquots were taken for UPLC-UV/MS analysis. Preparation of the tissue homogenates and UPLC-UV/MS parameters are given in supplementary information (Supplementary methods S3-S4). The half-live was calculated using the following equation:

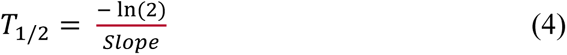

Where the slope is defined by linear regression assuming first order kinetics.

### Statistical analysis

Statistical analysis of PapRIV treated cells compared to vehicle treated cells was performed using one-way ANOVA; the Dunnett test was performed to adjust for multiple comparisons with the control group (placebo). A p-value of <0.05 was considered significant. The Mann-Whitney U test was used for the qPCR experiment. Linear and biphasic modelling was applied for the MTR experiments as described by Wynendaele *et al*. [14]. For the efflux and Caco-2 experiment, regular linear regression was applied. Data are expressed as mean ± SEM unless otherwise specified. Statistical analysis was performed and graphs were made using Graphpad Prism 6 (Graphpad Software, La Jolla, USA).

## Results

### PapRIV is able to reach the circulation and cross the blood-brain barrier

PapRIV showed a low transport rate across the Caco-2 monolayer with a P_app_ of 1.37 ± 0.21 x 10^-9^ cm/s and is thus potentially able to cross the intestinal wall and reach the circulation (Figure 1a). Indeed, PapRIV was detected in 4 out of 66 wild type mice plasma samples (Table 2 and Supplementary Fig. S1) which is an extra indication that the peptide is able to cross the gastro-intestinal tract and reach the circulation *in vivo*. Once circulating, the peptide is able to cross the blood-brain barrier based on experiments in the *in vivo* MTR mice model (Figure 1b). The pharmacokinetic parameters of PapRIV and of the negative and positive control, respectively BSA and Dermorphin, are given in Table 3. BSA and Dermorphin followed a linear model while the PapRIV peptide displayed a biphasic model with an initial steep influx followed by a steady state situation. PapRIV showed an initial brain influx rate (K_i_) of 6.95μl/(g x min) and can be classified as a peptide with a very high brain influx according to the classification system designed by Stalmans *et al*. [54]. Based on the capillary depletion experiment, it was observed that 87% of the peptide that is taken up by the brain eventually reached the parenchyma (Figure 1c and Table 3). Once the peptide enters the brain, no efflux back to the circulation is observed as the k_out_ was not significantly different from zero (Figure 1d and Table 3). In conclusion, the peptide is able to pass the Caco-2 monolayer, can be detected in mouse plasma and is able to pass the blood-brain barrier and reach the brain where possible effects can be exerted.

**Figure 1:**
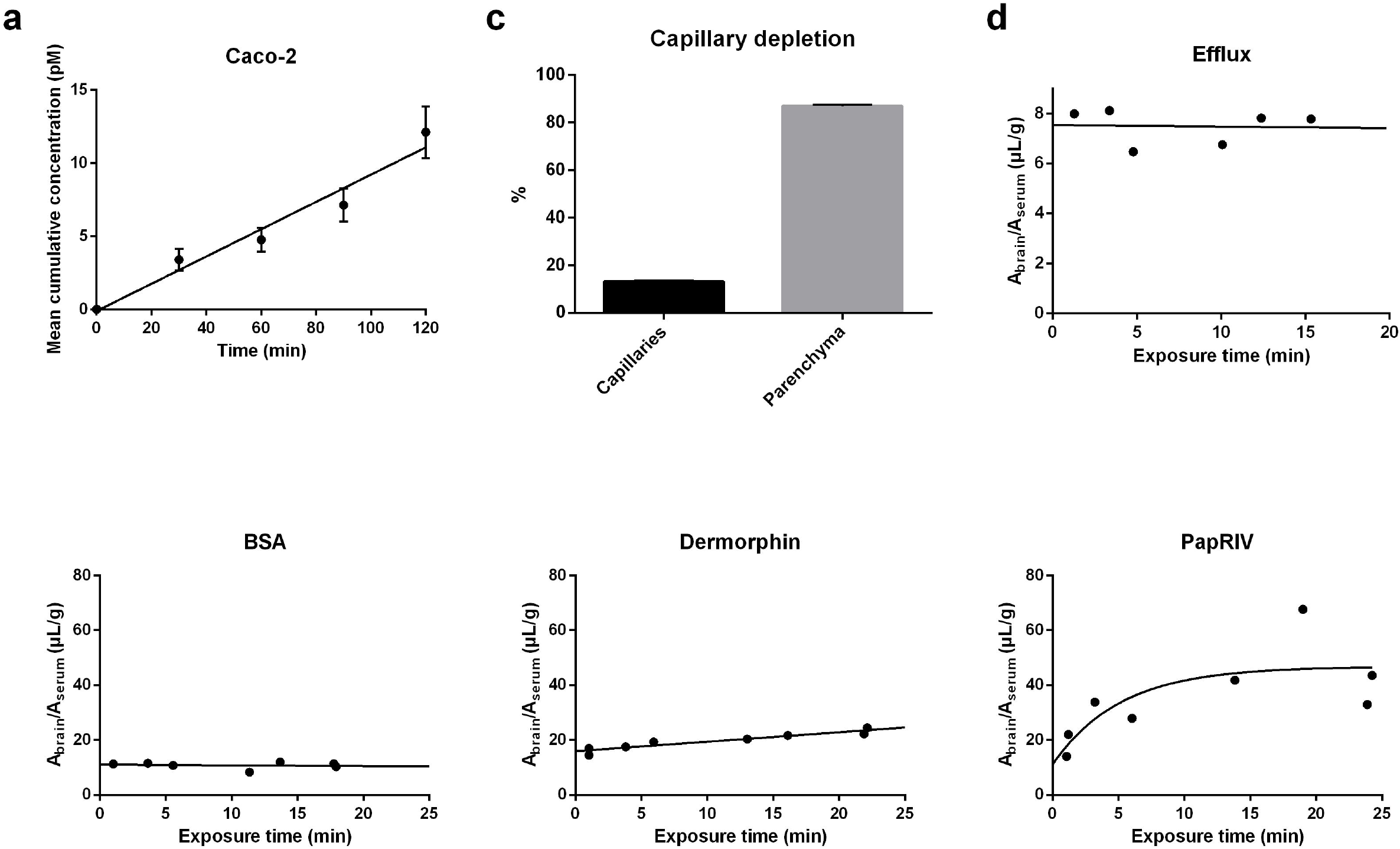
PapRIV passes the Caco-2 monolayer and blood-brain barrier. (a) The PapRIV peptide passes the Caco-2 monolayer and accumulates in the acceptor compartment (n = 6). (b) Multiple time regression analysis of BSA, Dermorphin and PapRIV across the blood-brain barrier. BSA and Dermorphin, the negative and positive controls respectively, follow a linear model while the PapRIV peptide is following a biphasic model with an initial steep influx followed by a steady state. (c) After 10 minutes, 87% of the peptide reaches the brain parenchyma while 13% remains in the capillaries (n = 2). (d) Efflux data of the peptide from the brain to the circulation.

**Table 2:**
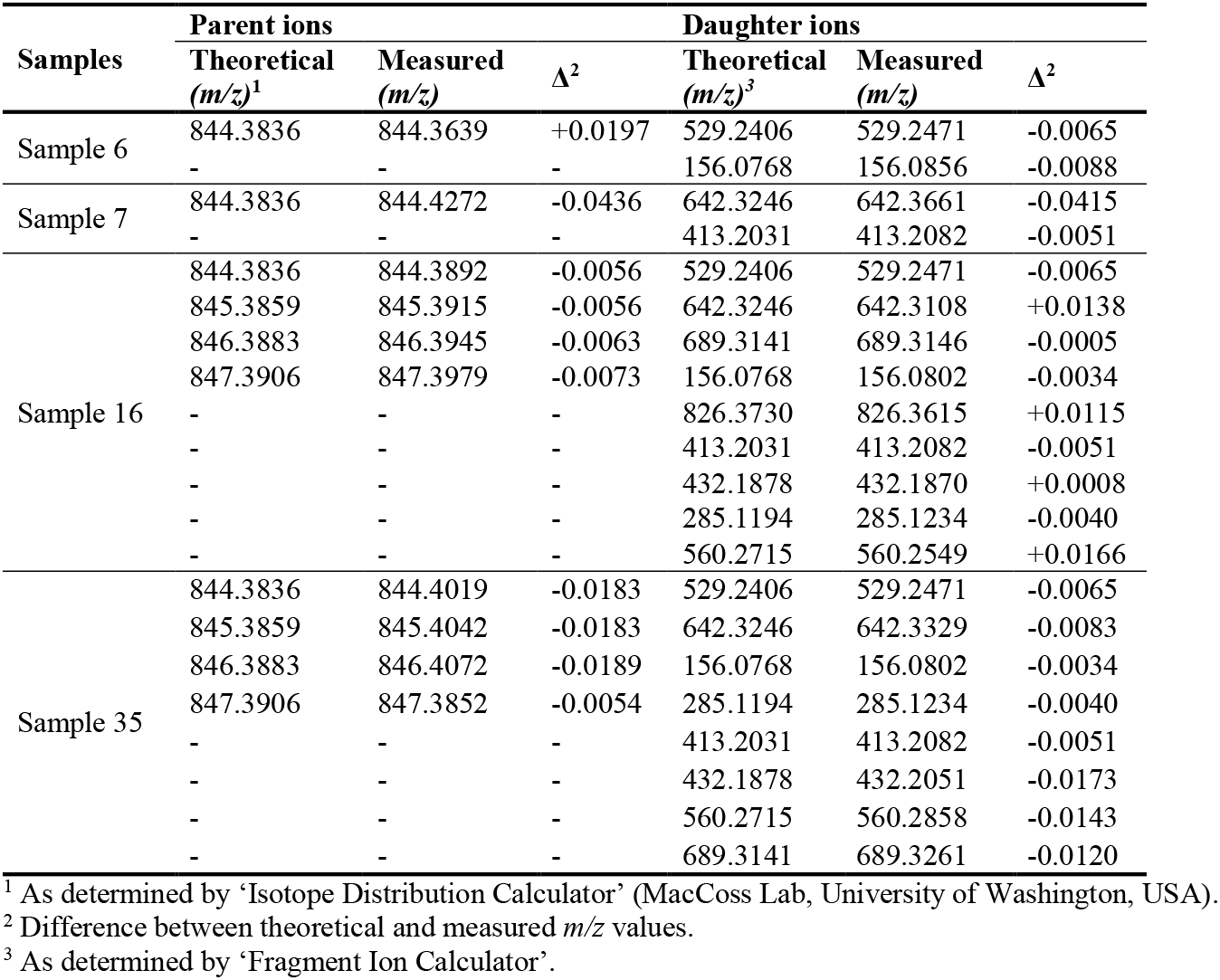
Measured parent- and daughter ions in positive mice plasma samples.

**Table 3:**
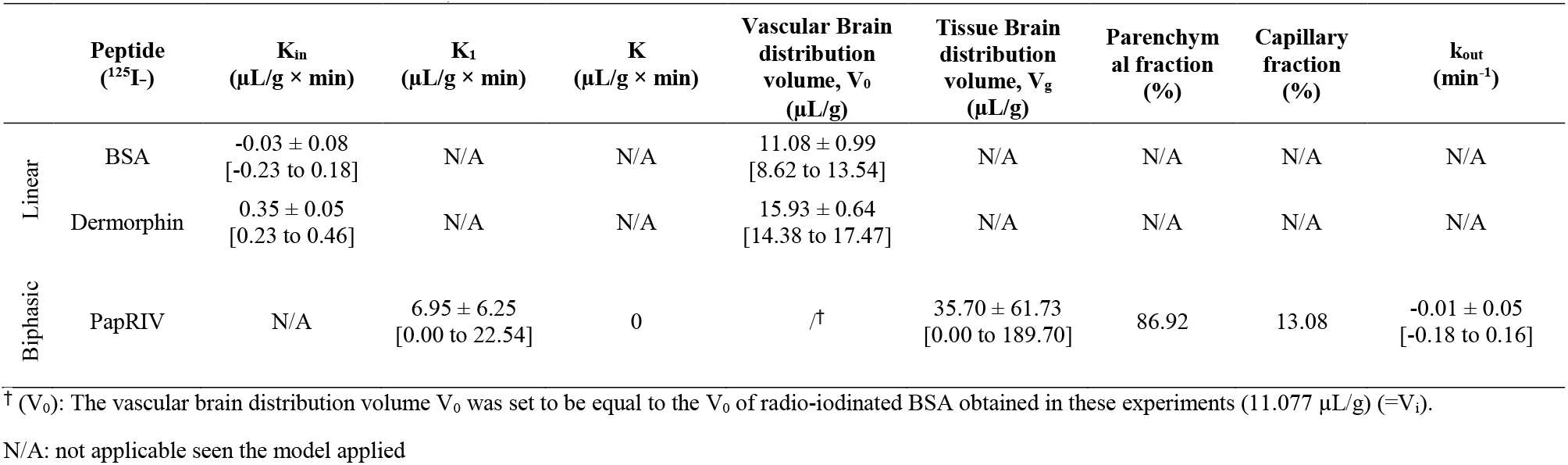
Overview of the multiple time regression results of BSA, Dermorphin and PapRIV using both the linear- and biphasic model (mean ± standard error, 95% CI interval between brackets)

|  | Peptide<br>(<sup>125</sup>I-) | K<sub>in</sub><br>(μL/g × min) | K<sub>1</sub><br>(μL/g × min) | K<br>(μL/g × min) | Vascular Brain<br>distribution<br>volume, V<sub>0</sub><br>(μL/g) | Tissue Brain<br>distribution<br>volume, V<sub>g</sub><br>(μL/g) | Parenchym<br>al fraction<br>(%) | Capillary<br>fraction<br>(%) | k<sub>out</sub><br>(min<sup>-1</sup>) |
| --- | --- | --- | --- | --- | --- | --- | --- | --- | --- |
| Linear | BSA | -0.03 ± 0.08<br>[-0.23 to 0.18] | N/A | N/A | 11.08 ± 0.99<br>[8.62 to 13.54] | N/A | N/A | N/A | N/A |
|  | Dermorphin | 0.35 ± 0.05<br>[0.23 to 0.46] | N/A | N/A | 15.93 ± 0.64<br>[14.38 to 17.47] | N/A | N/A | N/A | N/A |
| Biphasic | PapRIV | N/A | 6.95 ± 6.25<br>[0.00 to 22.54] | 0 | /† | 35.70 ± 61.73<br>[0.00 to 189.70] | 86.92 | 13.08 | -0.01 ± 0.05<br>[-0.18 to 0.16] |

† (V<sub>0</sub>): The vascular brain distribution volume V<sub>0</sub> was set to be equal to the V<sub>0</sub> of radio-iodinated BSA obtained in these experiments (11.077 μL/g) (=V<sub>i</sub>).
N/A: not applicable seen the model applied

### PapRIV shows pro-inflammatory effects on BV-2 microglia cells *in vitro*

The PapRIV peptide showed *in vitro* pro-inflammatory effects on the microglial BV-2 cell line, an immortalized murine microglial cell line which has proven its suitability for *in vitro* microglial research and investigation of neuro-inflammation [55–57]. The PapRIV peptide induced the production of the pro-inflammatory cytokines IL-6 and TNFα (Figure 2a); for IL-6, these changes were also confirmed at the mRNA level by qPCR (Figure 2b). This induction of pro-inflammatory cytokines was accompanied by an increase of intracellular ROS and an increased fraction of ameboid cells (Figure 2c-d, Supplementary Fig. S2). Treatment with 1 μg/mL LPS resulted in a fraction of ameboid cells of around ± 50% (data not shown). This microglial activation is mediated by an increased NF-kB nuclear translocation caused by decreasing IκBα levels, an inhibitory protein of NF-kB (Figure 2e-f). NIK expression was not observed in the cells (data not shown), thus indicating a canonical activation of the NF-kB pathway [58]. By synthesizing an alanine-scan of the sequence, we could identify the crucial amino acids of the peptide to exert its pro-inflammatory effects in the BV-2 microglial cells. When replacing aspartic acid or proline at respectively position 2 and 4 by an alanine residue, the corresponding peptide was not able anymore to induce IL-6 and TNFα production (Figure 2g). A scrambled control (DEHSFLP), which has the same amino acids as the native sequence but arranged in a random order, also showed no activating properties indicating that the specific sequence of amino acids is important for its function.

**Figure 2.**
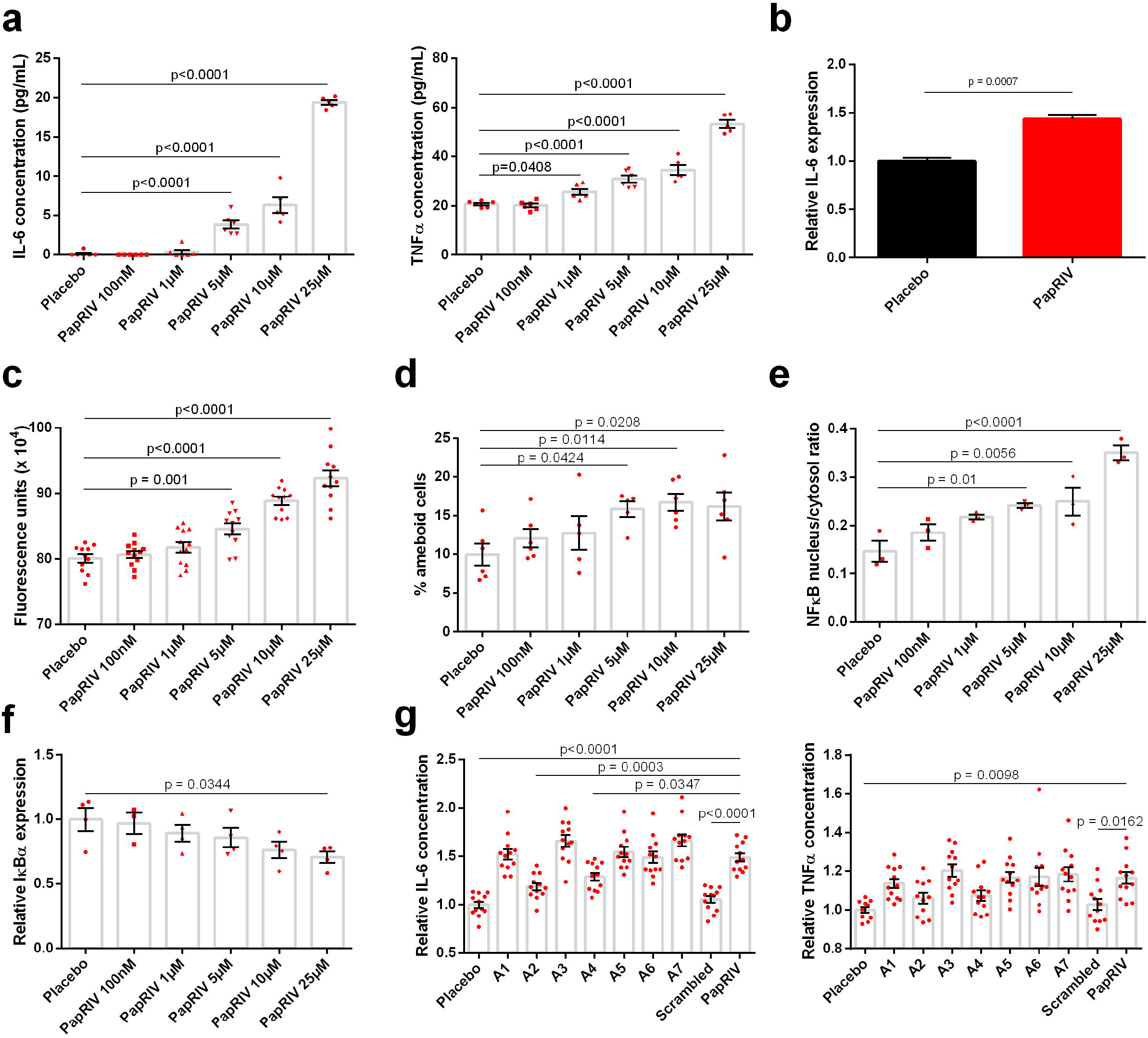
*In vitro* microglial activation of BV-2 cells by PapRIV. (a) IL-6 and TNFα levels increase after increasing concentrations of PapRIV (n = 6). (b) IL-6 mRNA expression is increased after treatment with 10 μM PapRIV (n = 6). *** = p<0.001 (Mann-Whitney U test) (c) Reactive oxygen species are formed after treatment (n = 12). (d) The fraction of ameboid cells, a marker for microglial activation, increased after treatment (n = 6). (e-f) Activation is mediated by an increased nuclear translocation of NF-kB caused by decreasing IκBα levels (n = 3). (g) Critical amino acids are identified by an alanine scan of the native sequence (10 μM, n = 12). Mean ± SEM, One-way ANOVA, post-hoc Dunnett.

### Conditioned medium of PapRIV treated BV-2 is toxic for SH-SY5Y neuroblast cells

Treatment of SH-SY5Y neuroblast cells with conditioned medium of BV-2 cells treated with PapRIV caused toxic effects on these neuroblast cells as demonstrated by a significant decreased viability of the cells (Figure 3a). This effect was not caused by direct actions of the peptide itself on the neuroblast cells as a direct treatment with the peptide caused no significant toxic effects (Figure 3b). The peptide thus shows indirect neurotoxic effect via microglia activation.

**Figure 3.**
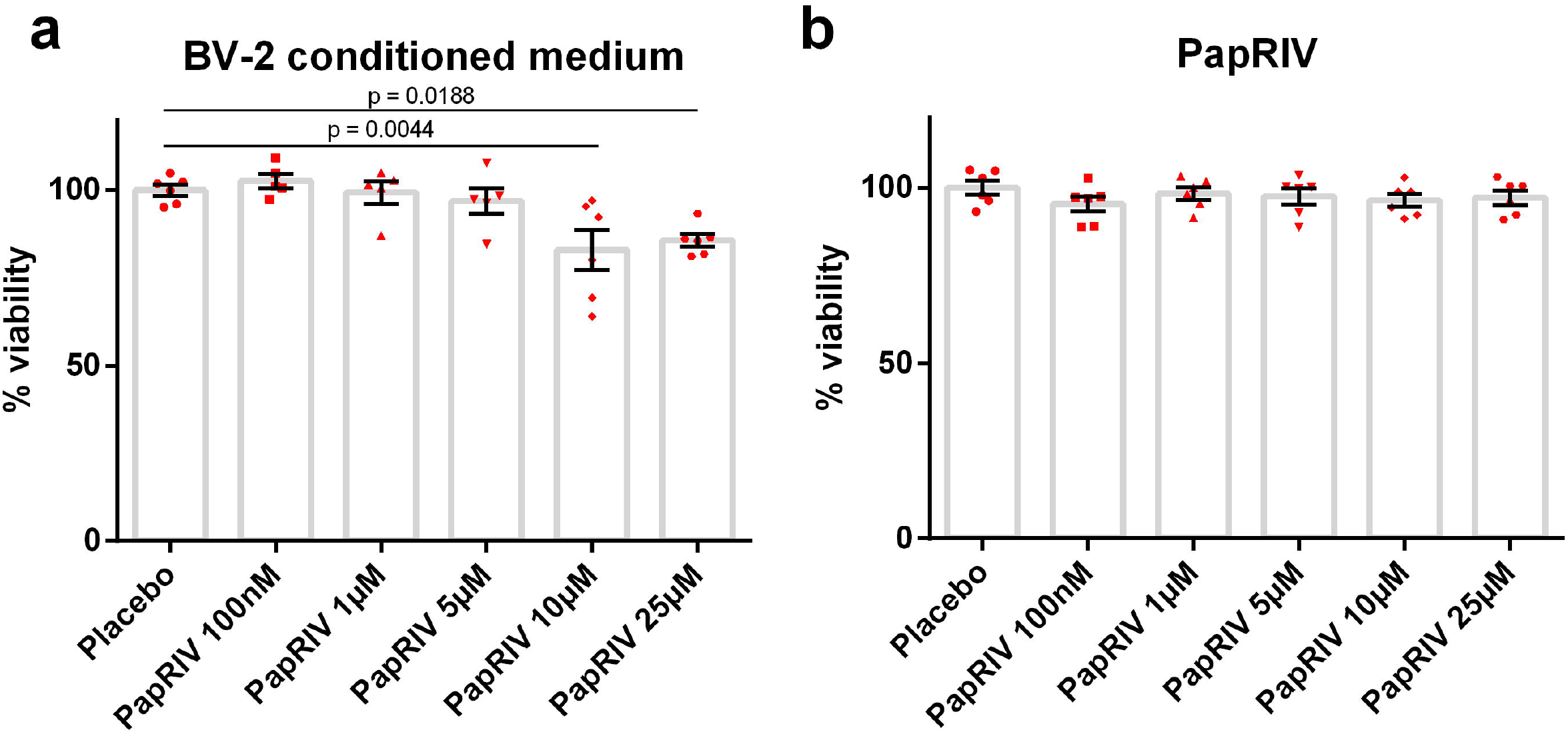
Indirect neurotoxic effects of PapRIV via microglia activation. (a) Treatment of SH-SY5Y cells with conditioned medium of PapRIV peptide-treated BV-2 cells is toxic for these neuroblasts (n = 6). (b) Direct treatment of SH-SY5Y cells with PapRIV shows no direct toxicity (n = 6). Mean ± SEM, one-way ANOVA, post-hoc Dunnett.

### Different metabolites of PapRIV are formed in serum, brain, liver, kidney and faeces

The metabolization rate of the peptide varied between different biological matrices such as serum, brain tissue, liver tissue, kidney tissue and faeces. The fastest metabolization rate was observed in kidney tissue, with a half-life of only 19.8 minutes. The half-lives in serum, faeces, liver and brain tissue were respectively 24.8, 89.7, 286 and 523 minutes while no metabolization was observed in colon tissue (Table 4, Supplementary Fig. S3). Different metabolites of the peptide could be identified. To exclude the possibility of non-enzymatic degradation, the chemical and protease-inactivated homogenate (by pre-heating the homogenate for 5 minutes at 95°C) stability of the peptide was determined. Except for kidney tissue, PapRIV remained stable in protease-inactivated homogenates, pointing an enzymatic degradation in serum, liver, brain and faeces (Supplementary Table S1). In protease-inactivated kidney homogenate, approximately 45% of peptide loss after 60 minutes can be explained by protein interaction with some kidney specific proteins. In serum, six different metabolites were formed. The MS-spectra of the different metabolites formed and the metabolic profiles in different tissues are given in supplementary information (Supplementary Fig. S4-S5). DLPFEH is the main metabolite which is formed in all matrices, metabolization of the native peptide to this metabolite is the main contributor of the low half-lives observed in kidney tissue and serum (Supplementary Fig. S5). Also in faeces, the matrix in which PapRIV is produced by gut bacteria, metabolization to DLPFEH occurred. Since this peptide still contains the two critical amino acids (D and P) at positions 2 and 4, respectively, the question was raised whether this peptide also showed microglial activating properties. As hypothesized, DLPFEH showed similar microglial activating properties as it was also able to induce IL-6 production to the same extent as the native sequence; the other metabolites showed no activity (Figure 4). Remarkably, SDLPF and DLPF showed no activating properties despite the presence of the two critical amino acids, thereby indicating that the presence of the C-terminal EH sequence is also necessary for the peptide’s action.

**Figure 4.**
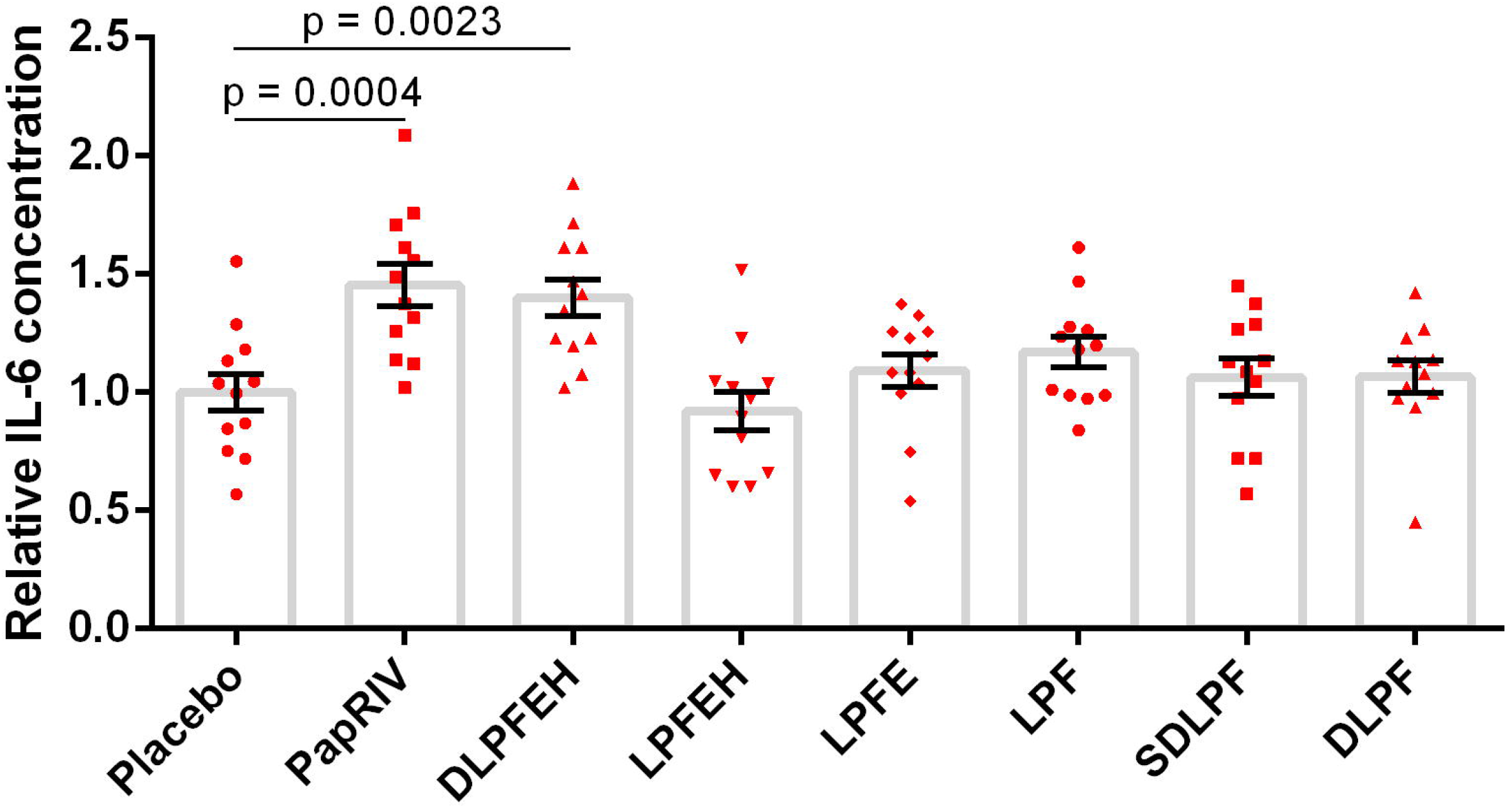
Activating properties of PapRIV metabolites on BV-2 microglia cells (10 μM, n = 12). Mean ± SEM, one-way ANOVA, post-hoc Dunnett.

**Table 4:** Peptide half-lives and formed metabolites in different biological matrices

| Matrix | T<sub>1/2</sub> (min) | Metabolites |
| --- | --- | --- |
| Serum | 24.8 | DLPFEH, LPFEH, LPFE, LPF, SDLPF, DLPF |
| Brain | 522.9 | DLPFEH, LPFEH |
| Liver | 285.7 | DLPFEH, LPFEH |
| Kidney | 19.8 | DLPFEH, LPFEH |
| Faeces | 89.7 | DLPFEH |
| Colon | - | - |

## Discussion

PapRIV, a QSP which is produced by *Bacillus cereus*, showed *in vitro* pro-inflammatory effects in BV-2 microglia cells. The peptide, which is mainly produced in the gut, is able to transfer across the gastro-intestinal tract and reach the circulation based on the *in vitro* Caco-2 model. A relatively low but appreciable P_app_ of 1.37 ± 0.21 x 10^-9^ cm/s is observed. A passive paracellular transport mechanism is suggested, as a comparable P_app_ was observed in the opposite basolateral-apical direction, as well as at 4°C (Supplementary Fig. S6). The low P_app_ can be explained by the low mass balance observed (<2%), indicating that peptide is lost during the experimental timeframe due to either cellular uptake or enzymatic degradation. Indeed numerous brush border membrane peptidases (i.a. aminopeptidases, dipeptidylpeptidase IV) are present on the apical side of the Caco-2 cells which may be responsible for enzymatic degradation of PapRIV during the experimental timeframe [59, 60]. The peptide was detected in 4 out of 66 mice plasma samples, while a protein BLAST search demonstrated that the sequence is not present in the mouse proteome and can thus not be the result of proteolytic cleavage of endogenous proteins. These findings supports the Caco-2 data that the peptide is able to cross the gastro-intestinal tract and reach the circulation *in vivo*. Presence of *B. cereus* in the human gastro-intestinal tract has already been demonstrated, up to 30% of vegetative cells and 100% of spores can survive gastric passage [61–64]. Gut permeability is also modulated by the intestinal microbiota. For example, expression of epithelial tight junction proteins is downregulated by several intestinal pathogens hereby increasing the permeability of the barrier [65]. The majority of *B. cereus* strains cause food poisoning and produce enterotoxins which are also able to increase vascular permeability [66, 67]. Spores are able to adhere in aggregates to both Caco-2 as small intestine gastro-intestinal epithelial cells which triggers germination and the production of enterotoxins; bacterial cells are thus in the proximity of the epithelial barrier facilitating the transport of metabolites [68]. Moreover, the intestinal barrier function is affected in several gastro-intestinal disorders such as IBD, IBS, celiac disease and obesity leading to a ‘leaky gut’ and an increased flux of luminal compounds towards the circulation [65]. Also in mental disorders such as ASD, ADHD and bipolar disorder, increased serum zonulin and/or claudin-5 levels are observed which is associated with respectively increased intestinal and BBB permeability [69, 70].

Once in the circulation, the peptide can reach the blood-brain barrier where it shows a very high brain influx according to the classification system of Stalmans *et al*. [54]. No significant brain efflux of the peptide was observed. PapRIV is thus able to reach the brain parenchyma where it can exert biological effects.

PapRIV showed *in vitro* microglia activating properties in BV-2 cells. A significant induction of both IL-6 and TNFα, two pro-inflammatory cytokines, in these cells is observed. These effects were accompanied by a significant increase of ROS. ROS are involved in the continued activation of microglia, even if the activating agent is already removed, a phenomenon called ‘reactive microgliosis’ [71]. This reactive microgliosis is explained by the presence of a self-amplifying loop: ROS are able to initiate NF-κB nuclear translocation and thus induction of gene transcription of pro-inflammatory mediators, while these pro-inflammatory cytokines consequently can increase ROS production [72]. Whether PapRIV has a direct effect on both factors or that one of both is indirectly affected by the other is yet unclear. In addition, microglia are extremely plastic and undergo a variety of spatio-temporal shape changes, dependent on their location and current role. Their morphology can range from highly branched, ramified cells with small cell bodies to ameboid, rounded cells with large cell bodies. When microglia are dormant, they have a lot of branches to survey the microenvironment; when they become activated and secrete pro-inflammatory cytokines, their morphology changes to ameboid by reorganizing proteins of the cell skeleton (actin, vimentin and microtubules) [73–75]. Here, an increased fraction of ameboid cells was observed after PapRIV treatment which is thus an additional proof of the microglia activating properties of this QSP. The effects are mediated by an NF-κB-dependent pathway, an increased nuclear translocation of NF-κB is observed which is caused by activation of the canonical pathway as a decrease of IκBα, a cytoplasmic inhibitory protein of NF-κB, and no NIK expression (data not shown) is seen [76]. When treating SH-SY5Y neuroblast cells with PapRIV conditioned BV-2 medium, toxic effects were seen at the higher PapRIV concentrations. Indeed, when treating SH-SY5Y cells with conditioned medium from activated BV-2 cells a decrease in viability is observed [77]. Several neurotoxic factors such as pro-inflammatory cytokines and extracellular ROS have been identified which may play a role in this microglia-mediated neurotoxicity [78]. Direct peptide treatment of the neuroblast cells did not result in a decreased viability, indicating that the toxic effects are mediated by the microglial activation. These results indicate the potential involvement of the peptide in microglia-mediated neurodegeneration.

To investigate the structure-activity relationship of the peptide, an alanine-scan was synthesized. By alternately replacing every amino acid with an alanine residue, crucial amino acids of the peptide can be identified. It is demonstrated that the second (aspartic acid) and the fourth (proline) amino acid are crucial for the peptide to exert its microglia activating effects.

A scrambled control peptide of the sequence did not show any activating effects; indicating that the specific peptide sequence is responsible for the pro-inflammatory actions.

Different metabolites of the peptide were *ex vivo* identified in different tissues with DLPFEH being the main metabolite which is formed in all tissues (serum, brain, liver, kidney and faeces). This metabolite contains the two critical amino acids and our experiment demonstrated that it remains active, thus contributing to PapRIV-mediated microglial activation. Two other metabolites, i.e. SDLPF and DLPF which are formed in serum, also contain the two critical amino acids, but are no longer active. This indicates that presence of the carboxy-terminal EH sequence is also necessary for the peptide’s actions. The half-live of the peptide varies from 20 minutes in kidney tissue to 523 minutes in brain tissue. In serum, a half-live of 25 minutes is observed. The differences observed between the tissues are due to the high concentration of proteases in serum and the renal brush border membrane, while in colon tissue, a much lower expression of brush border enzymes is observed [79]. Using PeptideCutter and PROSPER, an *in silico* prediction of the potential responsible proteases for the observed metabolites was made [80]. DLPFEH can be formed by cleavage of Asp-N endopeptidases at position 1 and DLPF due to an extra cleavage at position 5 by chymotrypsin or proteinase K. SDLPF can be formed by actions of chymotrypsin or proteinase K at position 5, while LPFE can be formed by the simultaneous action of formic acid at position 2 and glutamyl endopeptidase or matrix-metalloprotease 9 (MMP 9) at position 6. Finally, LPF can be formed directly by the combined actions of formic acid and proteinase K or chymotrypsin (Table 5).

**Table 5:**
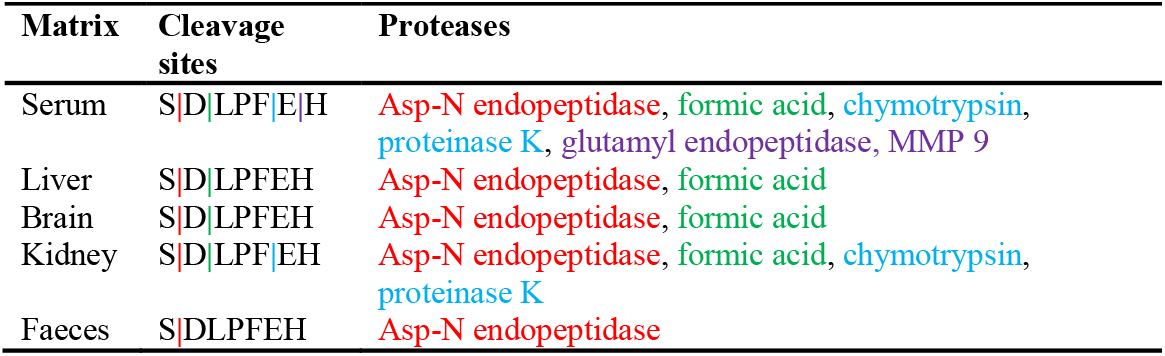
Overview of cleavage sites and *in silico* predicted responsible proteases

## Conclusion

PapRIV, a QSP produced by members of the *Bacillus cereus* group, shows *in vitro* activating properties towards BV-2 microglia cells. This QSP is produced in the gut and is able to cross the Caco-2 intestinal cell model via passive paracellular diffusion. Once reaching the circulation, it is able to cross the blood-brain barrier and reach the brain parenchyma. Presence of this QSP in mouse plasma was demonstrated for the first time. The peptide induces the expression of pro-inflammatory cytokines (IL-6 and TNFα) and ROS, and increases the fraction of ameboid cells via an NF-κB-dependent pathway in BV-2 microglia cells. Treatment of SH-SY5Y neuroblast cells with PapRIV conditioned BV-2 medium results in a decreased viability of these neuroblastoma cells, indicating indirect microglia-mediated neurotoxic effects of the peptide.

Overall, our *in vitro* obtained findings indicate for the first time a possible role of these bacterial quorum sensing peptides in gut-to-brain signaling, opening new avenues investigating their translational relevance.

## Acknowledgements

We thank Professor Alba Minelli for the kind gift of BV-2 cells.

## Authors’ contributions

YJ, ADS, ND and EW performed the experiments. MP and LF performed the synthesis of the alanine-scan and metabolites. AQ and CC performed the iPSC experiments. YJ, ND, ADS and BDS analyzed data. YJ, EW, DVD, PDD, PP, MBJ and BDS designed the experiments. YJ and BDS wrote the manuscript. The authors read and approved the final manuscript.

## Availability of data and material

The datasets used and/or analysed during the current study are available from the corresponding author on reasonable request.

## Ethics approval and consent to participate

Not applicable

## Consent for publication

Not applicable

## List of Abbreviations

QSP: Quorum sensing peptide
ICR-CD-1: Institute for Cancer Research, Caesarean Derived-1
BBB: Blood-brain barrier
ASD: Autism spectrum disorder
AD: Alzheimer’s disease
PD: Parkinson’s disease
ROS: Reactive oxygen species
FBS: Fetal bovine serum
NEAA: Non-essential amino acids
TMB: Tetramethylbenzidine
SEM: Standard error on mean
MMP-9: Matrix-metalloprotease 9

