## Supplementary information for "PapRIV, a BV-2 microglial cell activating quorum sensing peptide"

Supplementary information to: PapRIV, a BV-2 microglial cell activating quorum sensing peptide.

**Supplementary Methods**

**Supplementary method S1: Solid-phase peptide synthesis of alanine-scan, metabolites and scrambled control peptide**

**Synthesis of the alanine-scan peptides ADLPFEH, SALPFEH, SDAPFEH, SDLAFEH, SDLPAEH and SDLPFAH, and of the metabolites LPFEH and DLPFEH**

These peptides were synthesized manually by the solid-phase method following standard 9-fluorenylmethoxycarbonyl (Fmoc) chemistry. Fmoc-His(Trt)-Wang (150 mg, 0.23 mmol/g) was used as solid support. The resin was swollen with CH_2_Cl_2_ (1 × 20 min) and DMF (1 x 20 min). Peptide elongation was carried out through sequential Fmoc removal and coupling of the corresponding amino acids. Fmoc-Ala-OH, Fmoc-Ser(*t*Bu)-OH, Fmoc-Asp(O*t*Bu)-OH, Fmoc-Leu-OH, Fmoc-Pro-OH, Fmoc-Phe-OH and Fmoc-Glu(O*t*Bu)-OH were used as amino acid derivatives. Fmoc group removal was achieved with piperidine/DMF (3:7, 2 + 10 min). Couplings of the Fmoc-amino acids (4 equiv) were mediated by *N,N*-diisopropylcarbodiimide (DIC) (4 equiv) and ethyl cyanohydroxyiminoacetate (Oxyma) (4 equiv) in DMF at room temperature for 4 h under stirring. If the coupling was performed overnight, 3 equiv of each reagent were used. The completion of the reaction was checked with the ninhydrin [1] or chloranil [2] test. After each coupling and deprotection step, the resin was washed with DMF (6 × 1 min) and CH_2_Cl_2_ (3 × 1 min), and air-dried.

Once the synthesis of the peptide sequence was completed, the Fmoc group was removed and the peptidyl resin was treated with trifluoroacetic acid (TFA)/H_2_O/triisopropylsilane (TIS) (95:2.5:2.5) for 2 h. Following TFA evaporation and diethyl ether extraction, the peptide was dissolved in H_2_O and analyzed by HPLC. Finally, peptides were purified using a CombiFlash instrument, lyophilized, analyzed by HPLC and characterized by mass spectrometry.

**Synthesis of the alanine-scan peptide SDLPFEA and of the metabolites LPFE, SDLPF, DLPF and LPF**

These peptides were synthesized manually by the solid-phase method following standard Fmoc chemistry. MBHA resin (150mg, 0.5 mmol/g) was used as solid support. The resin was swollen with CH_2_Cl_2_ (1 × 20 min), DMF (1 × 20 min), piperidine/DMF (3:7, 1 × 5 min) and DMF (6 × 1 min). Then, the linker PAC (3 equiv) was coupled using DIC (3 equiv) and Oxyma (3 equiv) in DMF. The mixture was stirred overnight at room temperature. After this time, the resin was washed with DMF (6 × 1 min) and CH_2_Cl_2_ (3 x 1 min). The completion of the coupling was checked with the ninhydrin test. The conveniently protected C-terminus amino acid, Fmoc-Ala-OH, Fmoc-Glu(O*t*Bu)-OH or Fmoc-Phe-OH (5 equiv), was then coupled in presence of *N,N*-ethyldiisopropylamine (DIEA) (1 equiv), DIC (5 equiv) and 4-dimethylaminopyridine (DMAP) (0.5 equiv) in DMF under stirring for 2 h and the resin was washed with DMF (6 × 1 min) and CH_2_Cl_2_ (3 × 1 min). The coupling of this amino acid and the washes were performed twice. The loading of the resin was evaluated with the Fmoc test. Then, the resin was treated with acetic anhydride/pyridine/CH_2_Cl_2_ (7:7:86, 2 × 30 min) followed by washes with CH_2_Cl_2_ (3 × 2 min), DMF (3 × 2 min), MeOH (3 × 2 min), CH_2_Cl_2_ (2 × 2 min) and DMF (6 × 1min).

Peptide elongation was carried out through sequential Fmoc removal and coupling of the corresponding amino acids. Fmoc-Ser(*t*Bu)-OH, Fmoc-Asp(O*t*Bu)-OH, Fmoc-Leu-OH, Fmoc-Pro-OH, Fmoc-Phe-OH and Fmoc-Glu(O*t*Bu)-OH were used as amino acid derivatives. Fmoc group removal was achieved with piperidine/DMF (3:7, 2 + 10 min). Couplings of the Fmoc-amino acids (4 equiv) were mediated by DIC (4 equiv) and Oxyma (4 equiv) in DMF at room temperature for 4 h under stirring. If the coupling was performed overnight, 3 equiv of each reagent were used. The completion of the reaction was checked with the ninhydrin [1] or chloranil [2] test. After each coupling and deprotection step, the resin was washed with DMF (6 × 1 min) and CH_2_Cl_2_ (3 × 1 min), and air-dried.

Once the synthesis of the peptide sequence was completed, the Fmoc group was removed and the peptidyl resin was treated with TFA/H_2_O/TIS (95:2.5:2.5) for 2 h. Following TFA evaporation and diethyl ether extraction, the peptide was dissolved in H_2_O and analyzed by HPLC. Finally, peptides were purified using a CombiFlash instrument, lyophilized, analyzed by HPLC and characterized by mass spectrometry.

**Synthesis of the scrambled peptide DEHSFLP**

This peptide was synthesized manually by the solid-phase method following standard Fmoc chemistry as described for peptide SDLPFEA. After the coupling of the second amino acid, Fmoc-Leu-OH, the Fmoc group removal was achieved with tetrabutylammonium fluoride (TBAF)/ethanol/DMF (20 nM:2%:10 mL, 1 × 5 min) followed by washing the resin with DMF (6 × 10 sec) and CH_2_Cl_2_ (2 × 10 sec). The completion of the coupling was checked with the ninhydrin test [1]. Peptide elongation, cleavage and purification were then performed as described for peptide SDLPFEA.

**Supplementary method S2: UPLC-MS parameters Caco-2 assay**

Aliquots of the acceptor compartment are analyzed using the following validated UPLC-MS/MS method.

**Column** Acquity UPLC BEH C18 (2.1 × 100 mm, 1.7 µm) protected with a suitable guard column.

**Column temperature:** 60°C

**Sample temperature:** 10°C.

**Flow rate:** 0.5 mL/min

**Mobile phase:**

A: 93/2/5 (v/v) H_2_O + ACN + DMSO + 0.1% FA

B: 2/93/5 (v/v) H_2_O + ACN + DMSO + 0.1% FA

**Detection:**	MRM, the quantifier and collision energy are given in **Table 1.**

**Table 1: Quantifier of PapRIV.**

| **Peptide** | **Mother ion** | **Daughter ion** | **Collision energy MS** |
| --- | --- | --- | --- |
| PapRIV | 844.35 | 529.23 | 32 eV |

**Gradient composition:** The gradient program is presented in **Table 2**.

**Injection volume:** 10.0 µL

**Table 2: Gradient program**

| **Time (min)** | **Mobile phase A (%)** | **Mobile phase B (%)** |
| --- | --- | --- |
| 0.00 | 95 | 5 |
| 0.50 | 95 | 5 |
| 4.50 | 60 | 40 |
| 5.50 | 15 | 85 |
| 5.51 | 95 | 5 |
| 7.00 | 95 | 5 |

**Supplementary method S3: Tissue homogenate preparation**

**Preparation of Krebs-Henseleit buffer (pH 7.4) (KH-buffer)**

The powdered medium was dissolved in 900 mL water while stirring. To this solution, 0.3790 g CaCl_2_×2H_2_O and 2.098 g NaHCO_3_ was subsequently added while stirring. NaOH or HCl was used to adjust to pH 7.4. This solution was then further diluted to 1000 mL using ultrapure water.

**Preparation of tissue homogenates**

- The liver (n = 1), kidneys (n = 6), brains (n = 3), colons (n = 4) and faeces (0.196 g) of ICR-CD-1 mice were collected.
- After cleaning and rinsing the tissues using ice-cold KH buffer, the tissues were cut in little pieces and transferred into a 50 mL tube to which 36 mL ice-cold KH buffer was added.
- The tissues were then homogenized with at tissue homogenizer.
- After the larger particles were allowed to settle for about 30 minutes at 5°C, approximately 25 mL of the middle layer was transferred into a 50 mL Falcon tube using a plastic pipette.
- After shaking the homogenates, the homogenates were dispensed into 2 mL Eppendorf tubes, and stored at -35°C until use.
- The protein content of the homogenates were determined using the Pierce Modified Lowry Protein Assay method.
- Just before use, the homogenates were diluted to a protein concentration of 0.6 mg/mL (*i.e.* 300 µg / 500 µL) with KH buffer.

**Sample preparation**

- Into a 1.5 ml Protein LoBind tube, 900 µl of serum/tissue extract (500 µl of homogenate/serum and 400 µl of KH buffer, pH 7.4) was transferred.
- After temperature-equilibration at 37°C for 15 min while shaking at 750 rpm, 100 µl of peptide solution (0.1 mg/mL) was added followed by continued incubation at 37°C (while shaking), with 6×100 µl aliquots taken after 0, 5, 10, 30 and 60 minutes.
- The aliquots were immediately transferred into 500 µl Protein LoBind Eppendorf tubes containing 100 µl of 1% V/V trifluoroacetic acid solution in water, heated for 5 minutes at 95°C, and cooled for 30 minutes in an ice-bath.
- The samples were then centrifuged at 16,000 *g* for 5 minutes at 5°C.
- The supernatants was then analyzed by UPLC-UV/MS.

**Supplementary method S4: UPLC-UV/MS parameters *ex vivo* metabolization**

The analytical parameters of the UPLC-UV/MS method for the *ex vivo* metabolization are given in **Table 3.**

**Table 3: Analytical parameters of UPLC method**

| **Parameters** | **PapRIV** |
| --- | --- |
| Column | UPLC BEH-C18 (100 x 2.1 mm i.d.; 1.7µm) |
| Mobile phase A | H_2_O/ACN (95/5) + 0.1% FA |
| Mobile phase B | H_2_O/ACN (5/95) + 0.1% FA |
| Needle wash | H_2_O/ACN/DMSO (45/45/10), post injection (6 s) |
| Purge solvent | Methanol |
| Seal wash | Methanol/H_2_O (10/90) |
| Flow rate | 0.5 mL/min |
| Injection volume | 10 µL |
| Column temperature | 60 °C |
| Sample compartment | 10 °C |
| Detection | UV = 214 nm |
|  | MS = QDa |

The gradient program for the peptide is given in **Table 4**.

**Table 4: Gradient program**

| **Time** | **%A** | **%B** | **Curve** |
| --- | --- | --- | --- |
| 0 | 100 | 0 | 6 |
| 1 | 100 | 0 | 6 |
| 5 | 45 | 55 | 6 |
| 6 | 20 | 80 | 6 |
| 6.1 | 100 | 0 | 6 |
| 7 | 100 | 0 | 6 |

**Supplementary Figures**

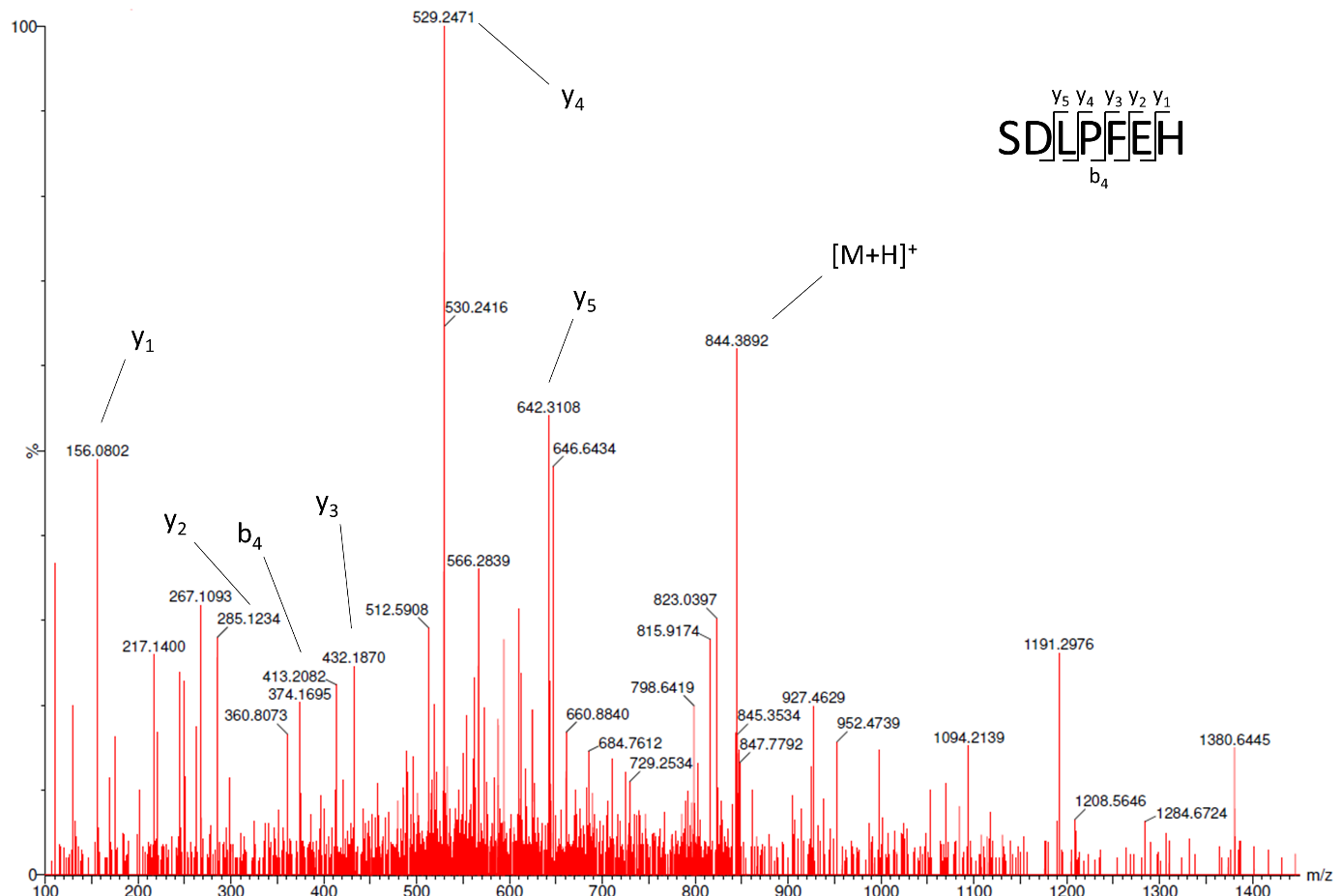

**Supplementary figure S1:** MS spectrum of a positive plasma sample obtained by the SYNAPT G2-Si system. The parent ion ([M+H]^+^) and detected y- and b-fragments of the parent ion (daughter ions) are indicated.

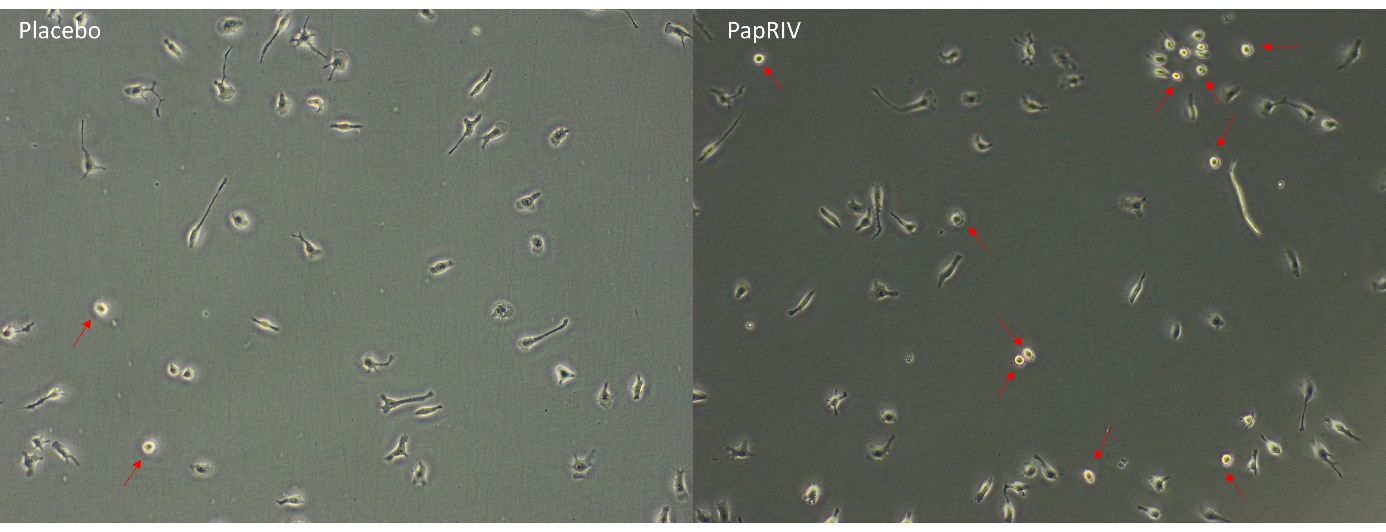

**Supplementary figure S2:** Morphology of placebo and PapRIV (25 µM) treated BV-2 cells. Red arrows indicate ameboid cells. The fraction of ameboid cells is increased after PapRIV treatment.

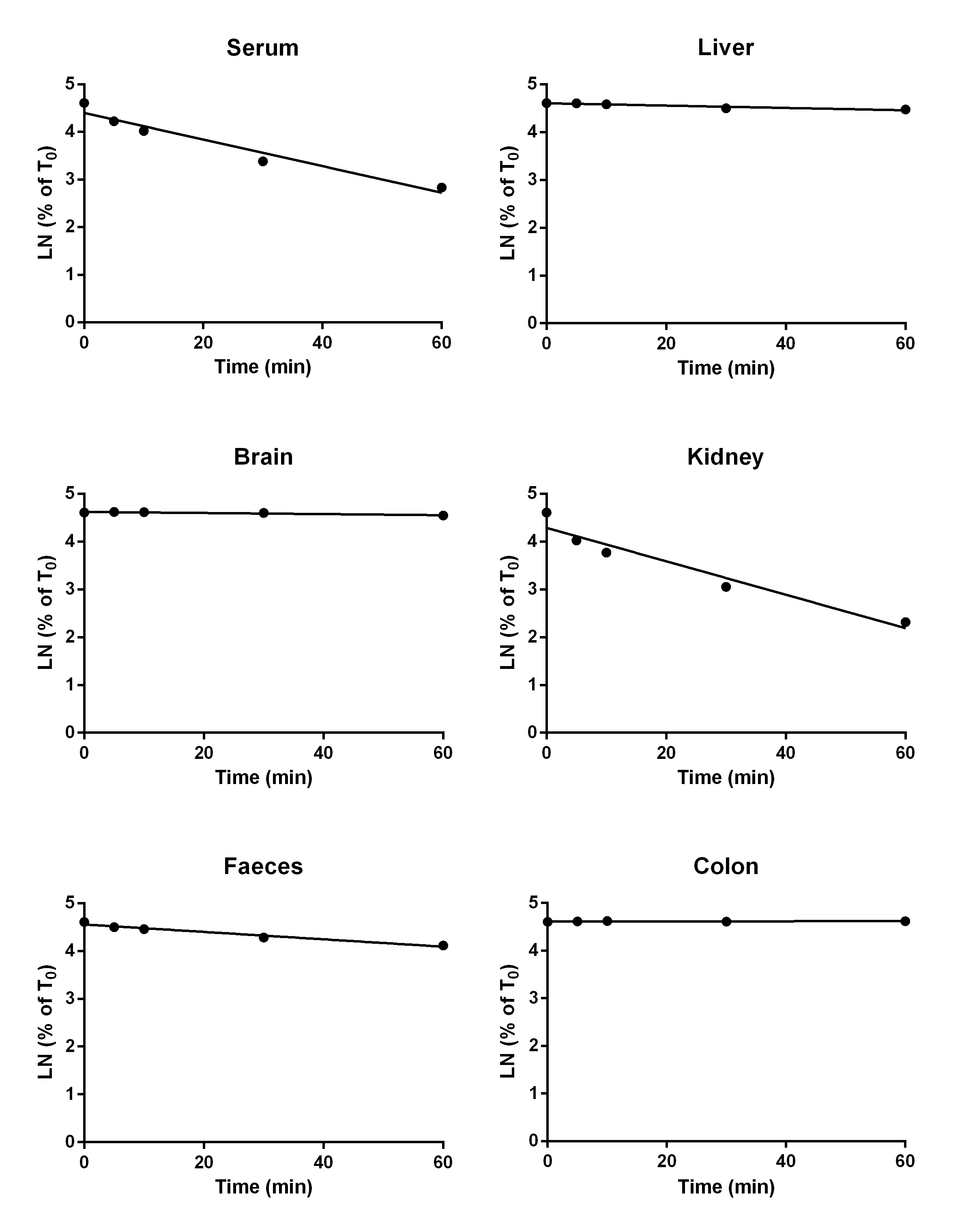

**Supplementary figure S3:** Degradation curves of PapRIV in different biological matrices (n = 1).

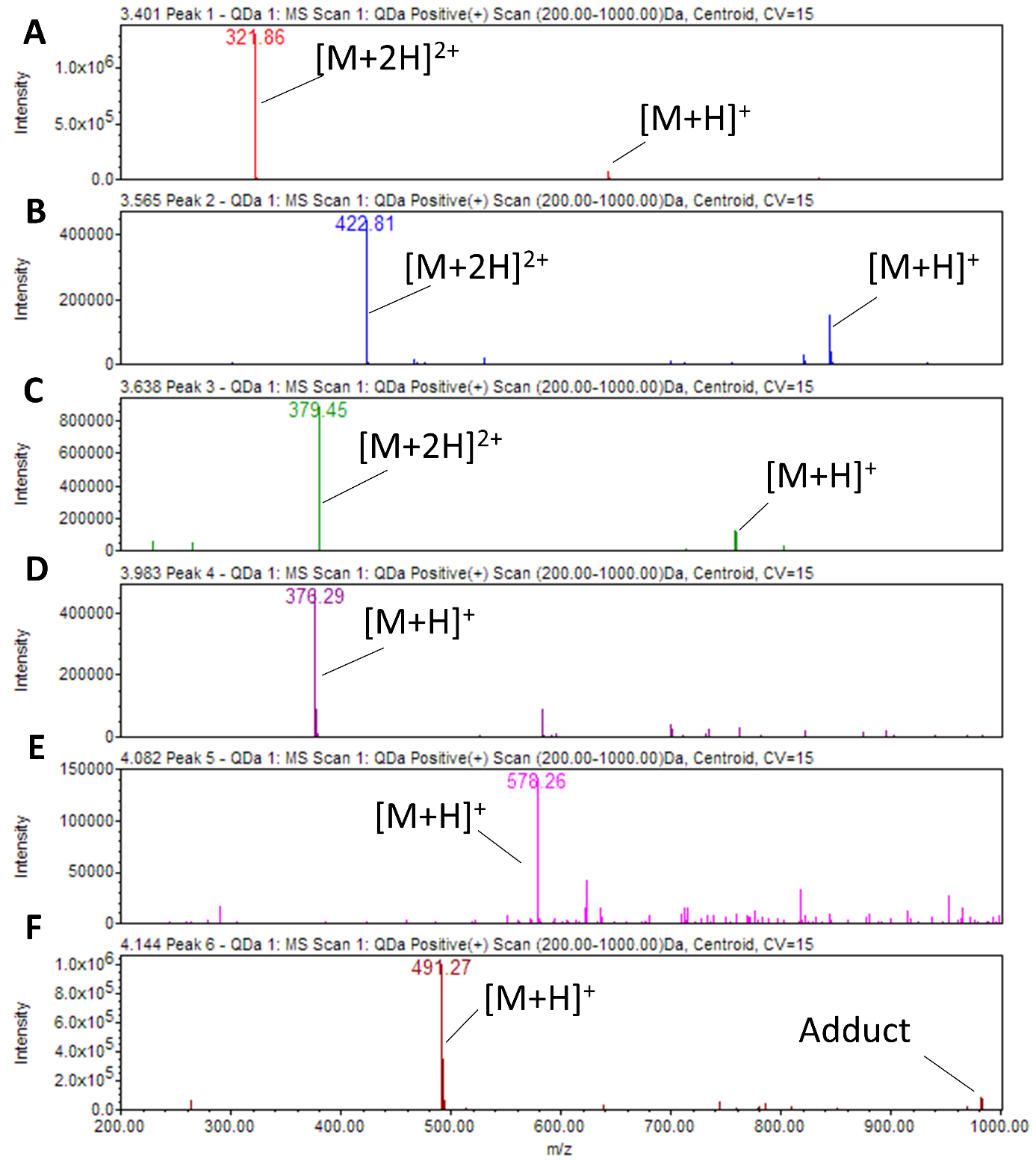

**Supplementary figure S4:** MS spectra of PapRIV and its metabolites: LPFEH (A), PapRIV (B), DLPFEH (C), LPF (D), SDLPF (E) and DLPF (F). LPFE was only visible in an extracted ion chromatogram (XIC).

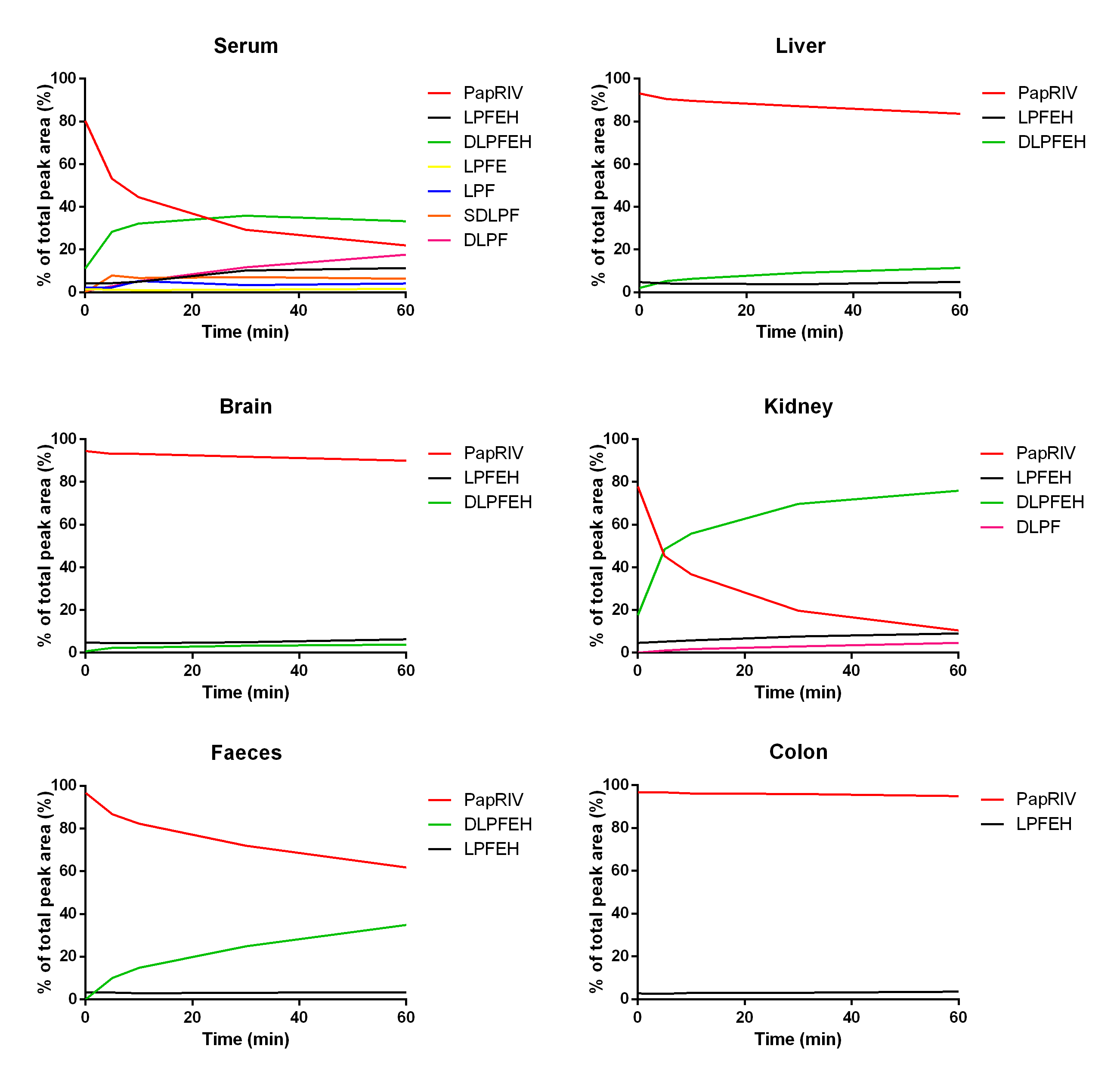

**Supplementary figure S5:** Metabolic profiles of PapRIV in different biological tissues. DLPFEH is identified as the main metabolite being the most abundant in all tissues.

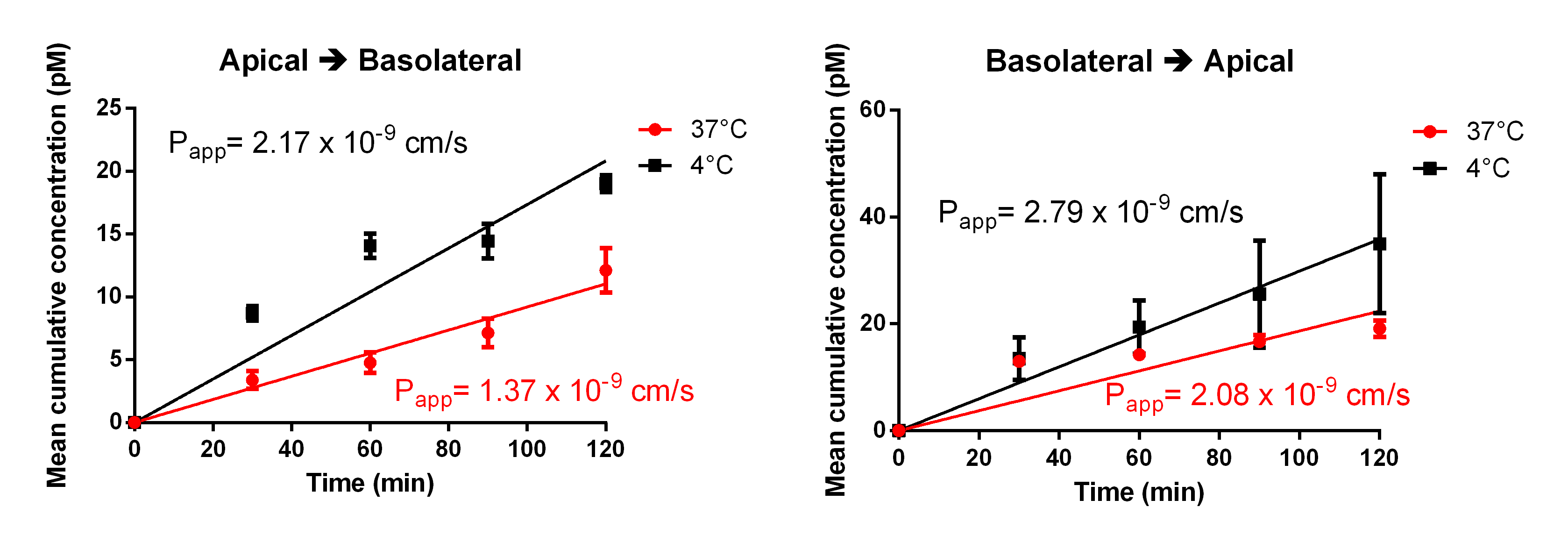

**Supplementary figure S6:** Apical to basolateral and basolateral to apical transport of PapRIV across the *in vitro* Caco-2 model at 37°C and 4°C (mean ± SEM, n = 6).

**Supplementary Tables**

**Supplementary table S1:** Chemical and protease-inactivated homogenate stability of PapRIV after 1 hour at 37°C.

| **Matrix** | **Remaining amount of PapRIV (%)** |
| --- | --- |
| KH-buffer | 99.8 |
| Inactivated serum | 99.0 |
| Inactivated brain | 101.0 |
| Inactivated liver | 101.6 |
| Inactivated kidney | 53.8 |
| Inactivated faeces | 101.4 |
| Inactivated colon | 99.2 |
